# simplePHENOTYPES: SIMulation of Pleiotropic, Linked and Epistatic PHENOTYPES

**DOI:** 10.1101/2020.01.11.902874

**Authors:** Samuel B. Fernandes, Alexander E. Lipka

## Abstract

**Motivation:** Advances in genotyping and phenotyping techniques have enabled the acquisition of a great amount of data. Consequently, there is an interest in multivariate statistical analyses that identify genomic regions likely to contain causal mutations affecting multiple traits (i.e., pleiotropy). As the demand for multivariate analyses increases, it is imperative that optimal tools are available to compare different implementations of these analyses. To facilitate the testing and validation of these multivariate approaches, we developed simplePHENOTYPES, an R package that simulates pleiotropy, partial pleiotropy, and spurious pleiotropy in a wide range of genetic architectures, including additive, dominance and epistatic models.

**Results:** We illustrate simplePHENOTYPES’ ability to simulate thousands of phenotypes in less than one minute. We then provide a vignette illustrating how to simulate a set of correlated traits in simplePHENOTYPES. Finally, we demonstrate the use of results from simplePHENOTYPES in a standard GWAS software, as well as the numerical equivalence of simulated phenotypes from simplePHENOTYPES and other packages with similar capabilities.

**Conclusions:** simplePHENOTYPES is a CRAN package that makes it possible to simulate multiple traits controlled by loci with varying degrees of pleiotropy. Its ability to interface with both commonly-used marker data formats and downstream quantitative genetics software and packages should facilitate a rigorous assessment of both existing and emerging statistical GWAS and GS approaches. simplePHENOTYPES is also available at https://github.coin/sainuelbfernandes/siinplePHENOTYPES.

## Background

The wealth of data available from high-throughput phenotyping platforms used in modern agronomical experiments is enabling unprecedented study into the genomic underpinnings of genotype-to-phenotype relationships [14, 12]. In particular, these data are making it possible to gain insight into the simultaneous contributions of genomic loci to multiple traits, a phenomenon known as pleiotropy [13]. To ensure that accurate inferences are being made from these data, the most appropriate statistical approaches must be used. One avenue towards assessing the performance of such approaches is to simulate correlated phenotypic data in which the pleiotropic and non-pleiotropic quantitative trait nucleotides (QTNs) underlying the genomic sources of phenotypic variability are known.

Several useful software packages have been developed to simulate correlated traits [9, 11, 4]. However, there is not a single package that enables the user to control for all possible parameters involved in the genetic architecture of complex traits. For instance, to the best of our knowledge, none of the currenlty availalble simulation packages allows the user to simulate spurious pleiotropy as defined by [13]. In this work, we present the R/CRAN package simplePHENOTYPES. This package uses real marker data to simulate additive (A), dominance (D), and additive x additive epistatic (E) QTN controlling multiple traits via pleiotropy, partial pleiotropy, and spurious pleiotropy. We maximize simplePHENOTYPES’ utility to the research community by ensuring its compatibility with popular data formats and state-of-the-art GWAS and GS software. Developmental versions, as well as vignettes, may be found at https://github.com/samuelbfernandes/simplePHENOTYPES.

## Implementation

simplePHENOTYPES is capable of simulating traits controlled by a wide range of genetic settings, as depicted in Figure 1. After reading in biallelic marker data in one of many formats (i.e., Numeric, HapMap, VCF, GDS, Plink Ped/Bed files) into R, the user specifies the desired genetic architecture using the *create_phenotypes()* function. In particular, the user has the option to indicate the degree of pleitoropy among the QTNs, the number of additive and non-additive QTNs, the degree of correlation between mutliple traits, and the heritability of each simulated trait.

**Figure 1:**
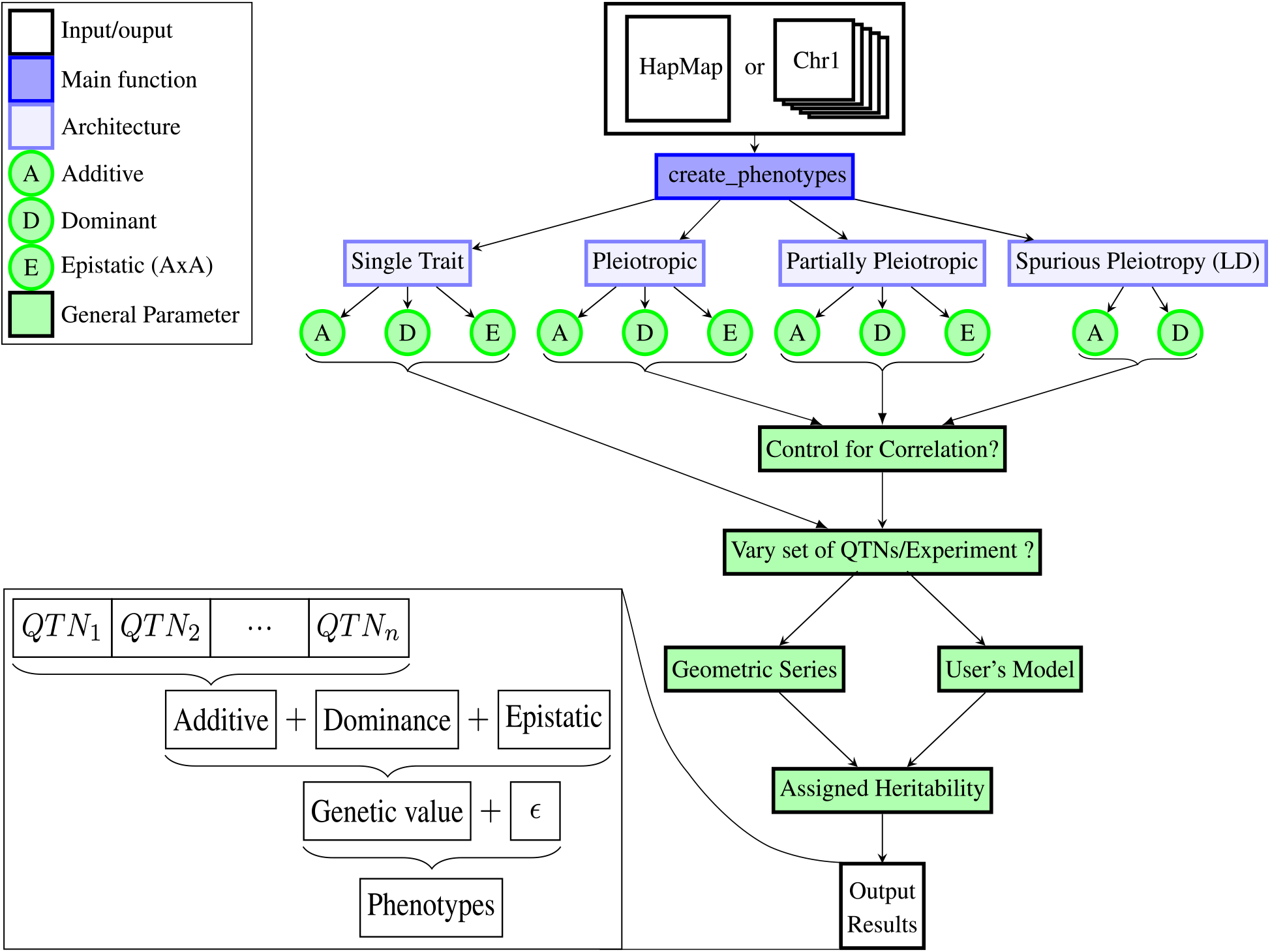
Workflow and main options implemented in simplePHENOTYPES to simulate single and multiple traits.

When simulating mutliple traits, the user has control over whether all QTNs control all traits (pleiotropy), a subset of the QTNs control all traits (partial pleiotropy), or separate QTNs in linkage disequilibrium (LD) control individual traits (spurious pleiotropy). To illustrate these three different scenarios, consider a set of 20 QTNs and two traits (*Trait j* and *Trait j’*), as depicted in Figure 2. In the pleiotropy scenario, the same 20 QTNs control both of the simulated traits. In the partial pleiotropy scenario, four QTNs control both traits, seven QTNs only control *Trait j,* and nine QTNs only control *Trait j*’. The final scenario, spurious pleiotropy, simulates two correlated traits where ten different QTNs control each trait. Here, simplePHENOTYPES randomly selects ten markers; each of these markers is in at maximum a user-specified amount of LD with one QTN controlling *Trait j*, as well as another QTN controlling *Trait j’*. The spurious pleiotropy depicted in Figure 2 is an implementation of what is defined in Figure 1-f Solovieff et al. [13], but simplePHENOTYPES also offers the option to simulate traits where the user-specified maximum LD refers to the maximum LD between each neighboring QTN pairs for the two traits. The user may select between the two LD options by using the *type_of_ld* input parameters of *create~phenotypes()* (see Listing 1 below).

**Figure 2:**
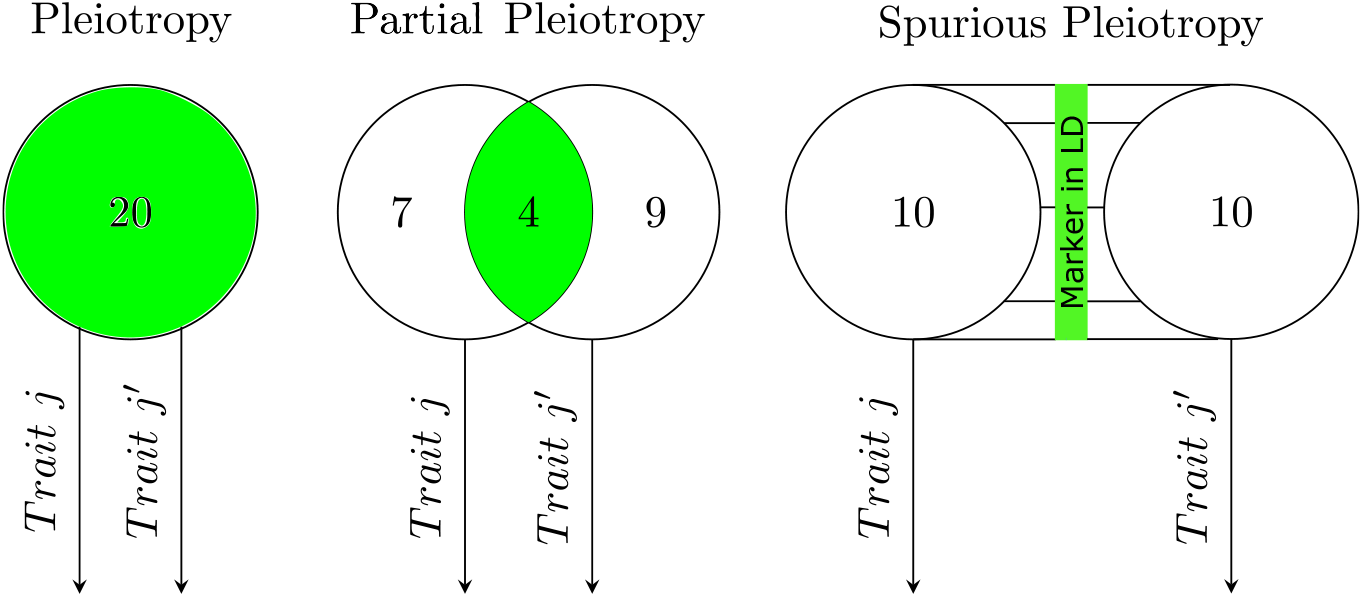
Three scenarios of multi-trait genetic architectures implemented in simplePHENOTYPES to simulate two traits *Trait j* and *Trait j’* controlled by a pool of 20 QTNs. Pleiotropic: 20 QTNs control both traits; Partially Pleiotropyc: 7, 4, and 9 QTNs controlling *Trait j,* both traits, and *Trait j’* only, respectively; Spurious Pleiotropy: 10 independent QTNs controlling each trait but in linkage disequilibrium (LD) with one SNP that is also in LD with QTNs from the other trait.

There is substantial flexibility for defining the number of additive and non-additive (i.e., dominance and additive x additive epistasis) QTNs controlling each trait, as well as their effect sizes. These effects may either be manually inputted in a list with a vector of effect sizes for each trait, or as a single value from which simplePHENOTYPES will create a geometric series of effect sizes [5]. The effect sizes will be assigned to each QTN, as described in Table 1. For a given number of QTNs and effect sizes, the user has the option to specify whether or not the same markers are to be the QTNs across all replicate traits through the *vary_QTN* input parameter of *create_phenotypes()*. The latter option of varying the markers randomly assigned to be QTN should allow the user to have a set of replicated traits where the QTN allele frequencies and the surrounding LD patterns differ for each simulated trait.

**Table 1:** Effect sizes assigned to each genotype.

| model |  | genotype |  |  |
| --- | --- | --- | --- | --- |
|  |  | -1 | 0 | 1 |
| additive | | $-a^*$ | 0 | $a$ |
| dominance | | 0 | $d^\ddagger$ | 0 |
| epistatic | -1 | $e^\dagger$ | 0 | $-e$ |
|  | 0 | 0 | 0 | 0 |
| | 1 | $-e$ | 0 | $e$ |

\*additive effect; ‡ dominance effect; † epistatic (additive x additive) effect.

It is possible to both indirectly and directly control the genetic correlation between traits simulated in simplePHENOTYPES. The genetic correlation between traits can be indirectly controlled by assigning different effect sizes to their shared QTNs. Thus, if the same two additive QTNs control two traits, denoted *Trait k* and *Trait k’*, with effect sizes 0.10 and 0.01 for *Trait k* and 0.40 and 0.16 for trait *Trait k’*, these two traits will have a genetic correlation 0 < *cor(Trait k,Trait k’)* < 1.

Alternatively, simplePHENOTYPES allows users to assign specific Pearson correlations between genetic values of the traits using a process known as whitening/coloring transformation [10]. To illustrate this process, let **Y** denote the scaled and centered genetic values (i.e., the cumulative values of the QTNs multiplied by their effect sizes) of each simulated trait. Let Σ denote the variance-covariance matrix of the simulated genetic values; since **Y** is scaled and centered, Σ is equal to a correlation matrix.

To modify **Y** so that the desired pairwise correlation between each genetic value is equal to those specified by the user in Σ’ (using the *cor* input parameter inside *create_phenotypes()),* the Cholesky decomposition is first applied to Σ. This phase, called the whitening transformation, calculates **X** = *L*^-1/2^**Y**, where *L* is a lower diagonal matrix defined from the Cholesky decomposition of Σ, i.e., Σ = *LL^T^*. Each of the genetic values in the resulting vector **X** are uncorrelated. To obtain a vector of genetic values **Y’** with the pairwise correlations specified in Σ’, a coloring transformation is applied to **X**, specifically **Y’** = *L*’^1/2^**X**. Similar to the whitening phase, *L’* is a lower diagonal matrix calculated from Σ’ = *L’L’^T^*, i.e. the Cholesky decomposition of Σ’. Finally, the process of centering and scaling the genetic values **Y’** is reversed so that these are back on the original scale.

Another feature of simplePHENOTYPES is to specify the heritabilities of each simulated trait. These heritabiltiies are subsequently used to simulate normally distributed random error terms that represent non-genetic sources of trait variability. By default, these residuals are independent, but the user may optionally specify a user-defined residual correlation *(cor_res*). Thus, for a given individual, the resulting simulated trait value is the sum of the simulated genetic values and this random error term.

We took several measures to ensure the quality, reproducibility, and accessibility of the results produced by simplePHENOTYPES. Upon the completion of simulating traits, simplePHENOTYPES will create a log file that will compare the estimated sample heritabilities and (when appropriate) genetic and residual correlations with those specified by the user, as well as confirm details on the specified genetic architecture. In addition, the variance explained by each QTN (calculated prior to the whitening/coloring transformation when the desired pairwise genetic correlations between traits are specified in Σ’), as well as new marker data without the SNPs selected to be QTNs, are optionally exported into the user’s specified working directory. Finally, the seed numbers used to select QTNs and for simulating the random non-genetic error terms for each simulated trait are saved as text files. These, along with genomic information on the markers selected to be QTNs, should facilitate the regeneration of the simulated traits whenever the need arises. The resulting simulated traits are saved in user-specified formats that are ready for downstream evaluation in external quantitative genetics software packages. Output compatibility includes GEMMA [15] and TASSEL [1]. Similarly, the possibility of saving the output as an R object ensures quick access by packages such as GAPIT [8] and rrBLUP [3].

## Results

simplePHENOTYPES is capable of simulating thousands experiments, i.e., single or multile phenotypes replicates, on a typical laptop computer in less than one minute (Figure 3). When we simulated phenotypes from a data set of 10, 650 markers genotyped on 280 maize lines from the Goodman-Buckler Diversity Panel [2, 6] on a single core of a 2.40 GHz MacBook with 64 GB of RAM, the median time to simulate 1,000 experiments (i.e., 1, 000 replicates of a trait with the same genetic architecture) for each evaluated scenario was 2.79 seconds (Figure 3). In particular, the median completion time for multivariate phenotypes was respectively 6.06, 15.57, and 10.79 seconds for the pleiotropy, partial pleiotropy, and spurious pleiotropy scenarios. Marker data sets in formats that require numericalization (e.g., HapMap formats) will require extra time for the numericalization step. The median time when this same data set was used in a HapMap format was 6.70 seconds (data not shown).

**Figure 3:**
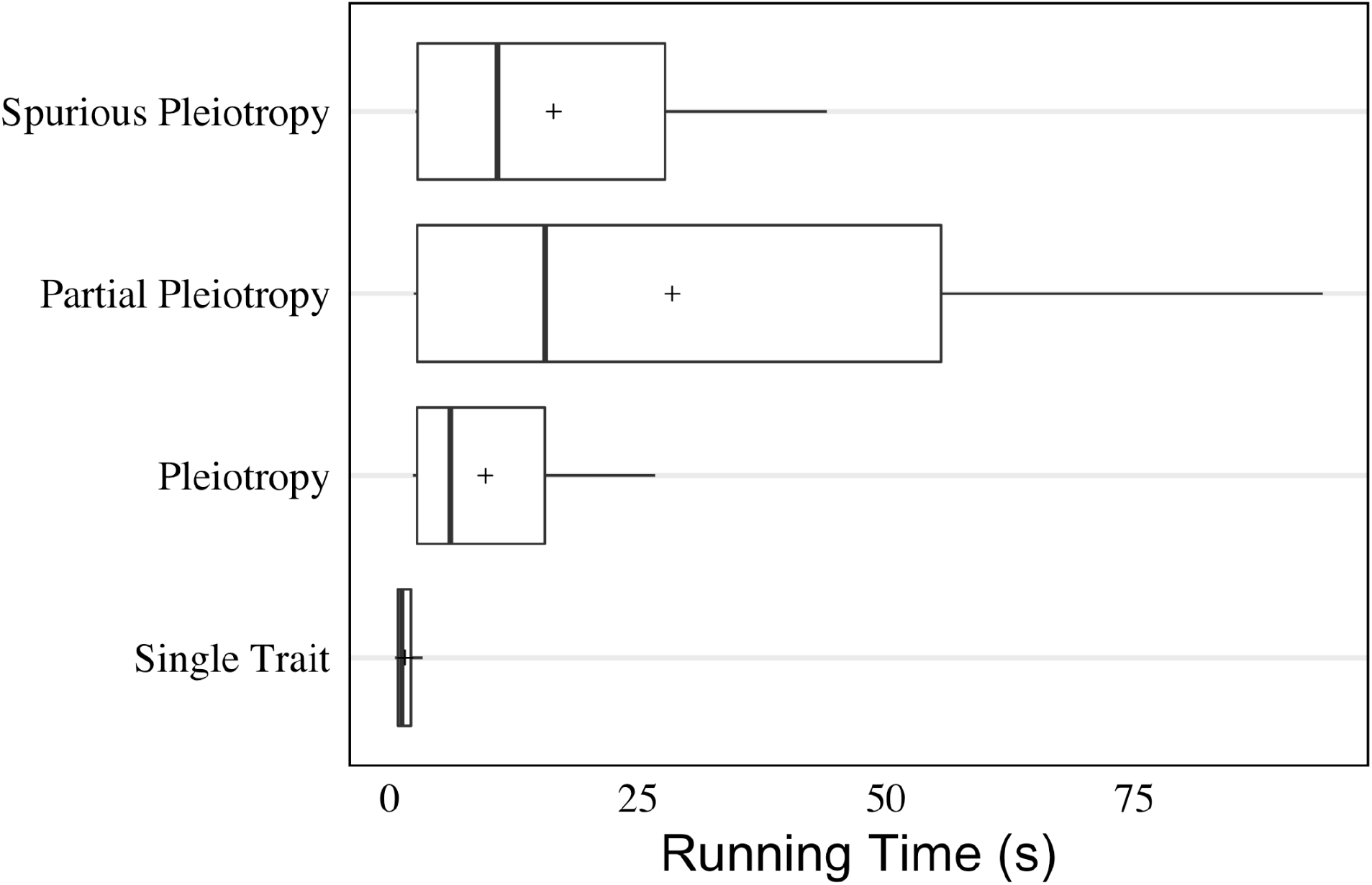
Boxplot of running time, in seconds, required by simplePHENOTYPES to simulate 1,000 replicate phenotypes under single trait, pleiotropic, partial pleiotropy, and spurious pleiotropy architecture. In the latter three scenarios, two traits were simulated for each replicate phenotype. This simulation was conducted on a single core of a 2.4 GHz MacBook with 64 GB of RAM. The data set used contained 10,650 SNPs and 280 maize individuals in numeric format. The symbol “ + “ denotes the mean of each architecture simulated.

To illustrate the facility of simulating phenotypes consisting of multiple traits in simplePHENOTYPES, we present a sample R script for simulating two traits from the spurious pleiotropy scenario (Listing 1). Here, the same data set of 280 maize lines form the Goodman-Buckler diversity panel is used to simulate two traits controlled by three additive QTNs (*addjQTN_num* = 3). Both traits have a heritability of 0.5. The additive effect sizes for the three QTNs of trait 1 (trait 2) are 0.2 (0.3), 0.1 (0.2), and 0.05 (0.1). The LD between a given QTN of each trait and the corresponding common SNP (*type_of_ld = “indirect”*) was at maximum 0.7 (*Id* = 0.7).

**Listing 1:**
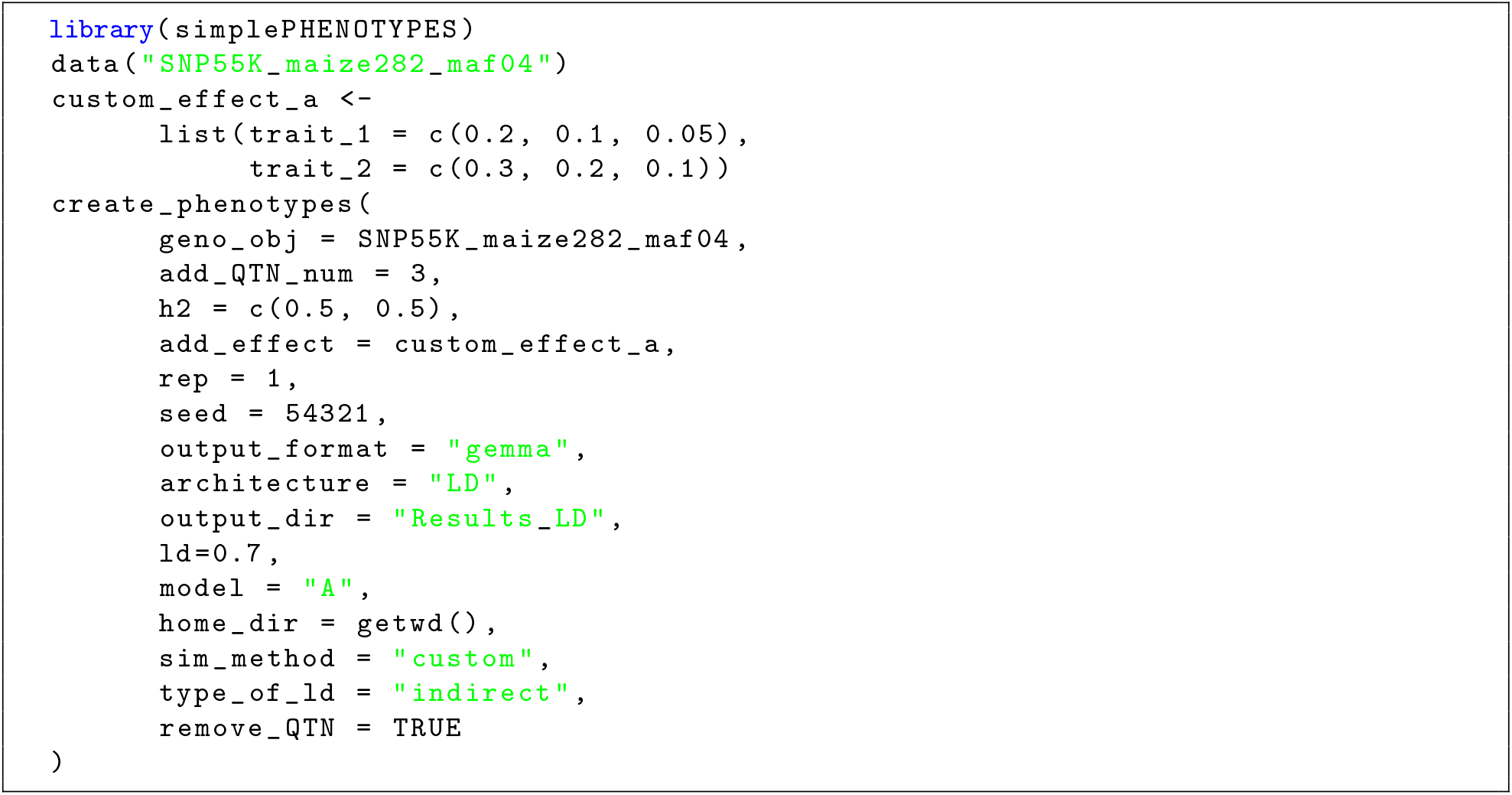
Example of a script for simulating two traits controled by spurious pleiotropy

Detailed information on the SNPs selected to be QTNs and a summary of their LD are presented at Tables 2 and Table 3, respectively. To illustrate the use of these simulated traits in downstream quantitaitve genetics analysis, we removed the SNPs selected to be QTNs from the marker data set *(remove~QTN* = *TRUE*) and conducted a multi-trait GWAS using GEMMA [16] (Figure 4). Similar GWAS examples for phenotypes simulated under the pleiotropy (Tables S1, S2, and Figure S1) and partial pleiotropy scenarios (Tables S3, S4, and Figure S2) are presented in the section 2 of the Supplementary File. As expected, in all scenarios the peak-assocaited markers from GWAS were located in the vicnity of the largest-effect QTNs.

**Figure 4:**
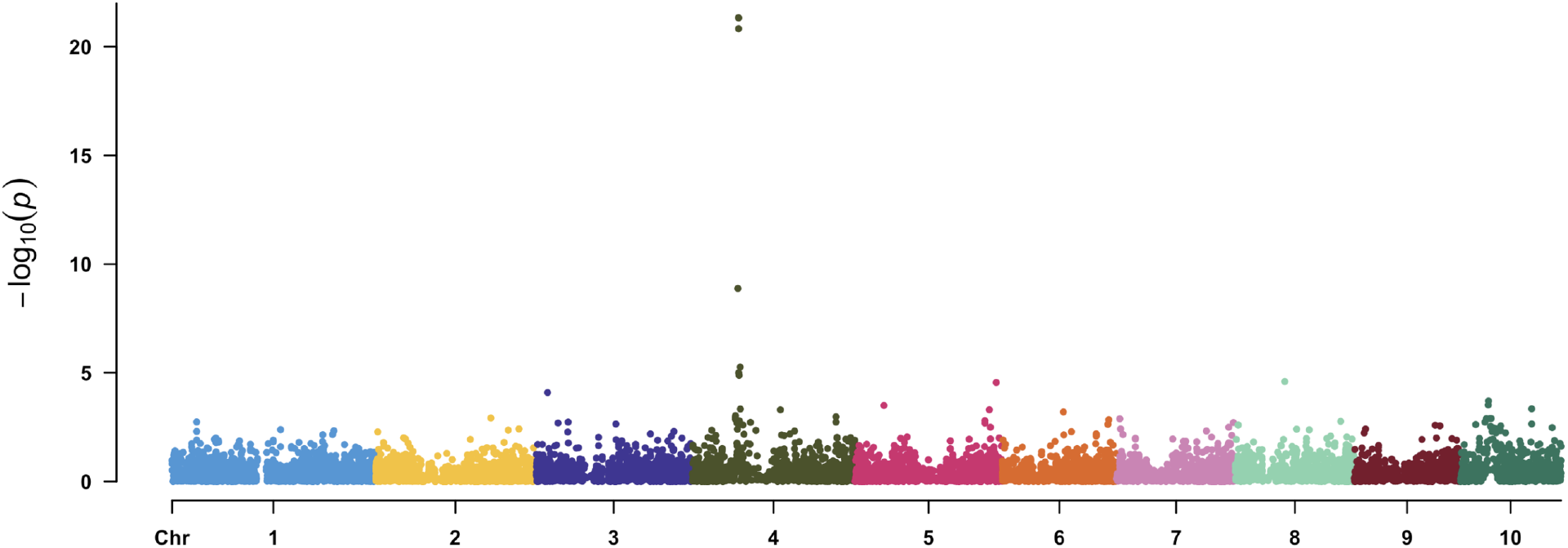
Manhattan plot of a multi-trait GWAS conducted on two phenotypes simulated under the spurious pleiotropy genetic architecture.

**Table 2:**
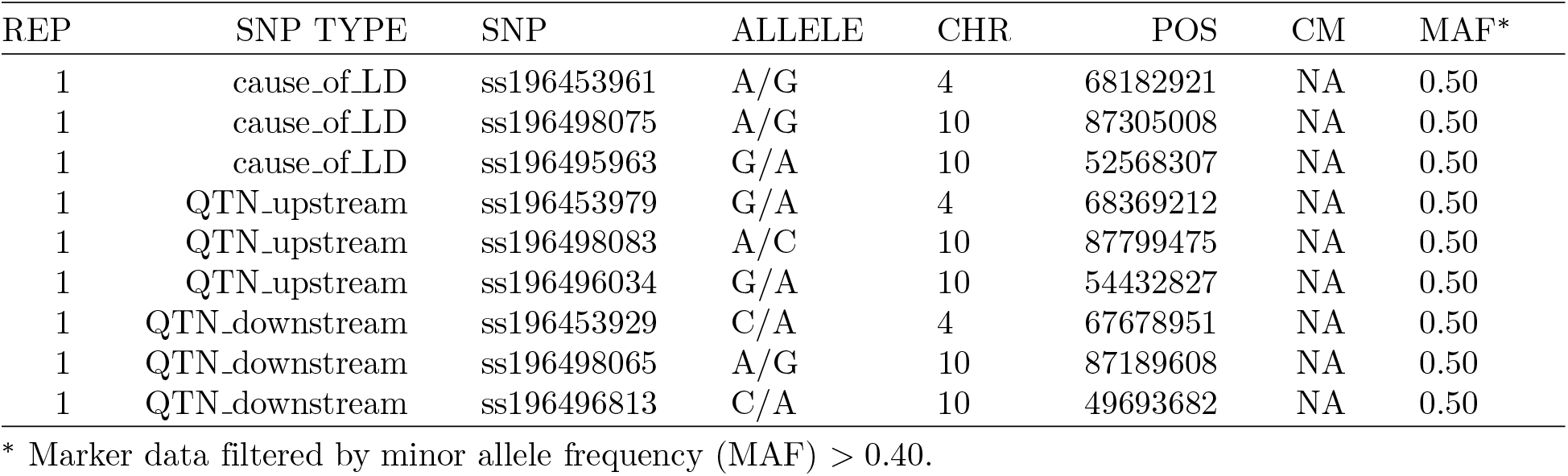
Map and minor allele frequency information of SNPs randomly selected to be used as additive quantitative trait nucleotide in each replication.

**Table 3:** Summary linkage disequilibrium information on quantitative trait nucleotide of different traits.

| REP | SNP causing LD | input LD (absolute value) | Actual LD with QTN of Trait 1 | Actual LD with QTN of Trait 2 | QTN for trait 1 | QTN for trait 2 | LD between QTNs |
| --- | --- | --- | --- | --- | --- | --- | --- |
| 1 | ss196453961 | 0.70 | 0.16 | 0.11 | ss196453929 | ss196453979 | 0.11 |
| 1 | ss196498075 | 0.70 | 0.24 | 0.45 | ss196498065 | ss196498083 | -0.35 |
| 1 | ss196495963 | 0.70 | 0.66 | 0.62 | ss196496813 | ss196496034 | 0.62 |

To demonstrate equivalence of results between simplePHHENOTYPES and similar packages, we compared phenotypes simulated by simplePHENOTYPES to those from AlphaSimR [4], SimPhe [7] and PhenotypeSimulator [9] when equivalent genetic architectures were specified. Due to differences between simplePHENOTYPES and PhenotypeSimulator concerning specific input parameters and implementations used to simulate phenotypes, we compared the ability of both packages to simulate phenotypes controlled by only additive QTNs. Additionally, we compared simplePHENOTYPES to AlphaSimR and SimPhe for phenotypes controlled by additive, dominance, and additive x additive epistatic QTNs. We showed that when phenotypes with the same QTNs and effect sizes where simulated under a heritability of 1, the simulated phenotypes from simplePHENOTYPES were identical to the other software pakcages (please see the Supplementary File and vignettes at the GitHub page for more details and sample code).

## Conclusions

simplePHENOTYPES makes it possible to simulate multiple traits controlled by loci with varying degrees of pleiotropy. Its ability to interface with both commonly-used marker data formats and downstream quantitative genetics software and packages should facilitate the rigorous assessment of both existing and emerging statistical GWAS and GS approaches.

## Supplementary information

Supplementary information accompanies this paper at: https://github.com/samuelbfernandes/simplePHENOTYPES/blob/master/vignettes/Supplementary.pd

## Acknowledgements

We acknowledge Candice Hirsch, Rafael Della Coletta, Brian Rice and Kaio Olimpio Das Graças Dias for their independent testing and suggestions that improved the usability of simplePHENOTYPES.

## Availability and requirements

Project Name: simplePHENOTYPES

Project Home page: https://github.com/samuelbfernandes/simplePHENOTYPES

Operating system: Tested on Linux, Windows and macOS

Programming languages: R

Other requirements: R 3.5 or higher

License: MIT license

Any restrictions to use by non-academics: No (free software)

## Author’s contributions

SBF led the development and implementation of simplePHENOTYPES and wrote the manuscript, while AEL supervised the development of simplePHENOTYPES and wrote the manuscript.

## Ethics approval and consent to participate

Not applicable.

## Consent for publication

Not applicable.

## Author details

^1^Department of Crop Sciences, University of Illinois at Urbana-Champaign, Urbana, IL, 61801, USA

