## Supplementary File for "simplePHENOTYPES: SIMulation of Pleiotropic, Linked and Epistatic PHENOTYPES"

This is the supplementary file that includes figures and performance assessments for Fernandes and Lipka (2020). All the examples below were run on simplePHENOTYPES v1.2.4.

### 1 Benchmarking against other packages

In this performance assessment, we benchmarked simplePHENOTYPES against AlphaSimR (Faux et al., 2016), SimPhe (Jiang and Pütz, 2018), and PhenotypeSimulator (Meyer and Birney, 2018). For all benchmarking, we used a data set composed of 100 individuals and two SNPs simulated with AlphaSimR. Additionally, in all scenarios, the simulated heritability was 1, and the allelic effects were the same between simplePHENOTYPES and the package into consideration. To compare simplePHENOTYPES to AlphaSimR and SimPhe, we simulated two traits under additive (A), dominance (D), and additive x additive epistatic (E) models, controlled by two, two, and one quantitative trait nucleotide (QTN), respectively. The comparison with PhenotypeSimulator, on the other hand, involved two additive traits controlled by two QTNs. Please note that rather than showing the superiority of a given package, the goal of these comparisons is to show that some of the genetic models implemented are equivalent.

```
#devtools::install_github("samuelbfernandes/simplePHENOTYPES")
library(simplePHENOTYPES)
library(PhenotypeSimulator)
library(SimPhe)
library(AlphaSimR)
```

#### 1.1 simplePHENOTYPES vs AlphaSimR

##### 1.1.1 AlphaSimR ADE

Because AlphaSimR samples the allelic effects from a normal or gamma distribution, we used those allelic effects as the user-defined allelic effects for simplePHENOTYPES. After the addition of the intercept simulated by AlphaSimR, both packages produced the same phenotypes for both traits.

- heritability = 1
- number of traits = 2
- genetic model = A, D, and E
- Number of QTNs: A = 2, D = 2, E = 1
- allelic effects = N($\mu$ = 1, $\sigma^2$ = 1)

```r
set.seed(1)
founderPop = quickHaplo(nInd = 100,
                        nChr = 2,
                        segSites = 1)
SP_ade = SimParam$new(founderPop)
SP_ade$addTraitADE(
  nQtlPerChr = 1,
  mean = c(1, 1),
  var = c(1, 1),
  meanDD = c(1, 1),
  varDD = c(1, 1),
  relAA = 1)
SP_ade$setVarE(H2 = c(1, 1))
pop = newPop(founderPop, simParam = SP_ade)
pheno_ade_alpha <- pheno(pop)
gv <- genParam(pop, simParam = SP_ade)
QTNs_ade <- pullQtlGeno(pop, trait = 1, simParam = SP_ade)
```

##### 1.1.2 simplePHENOTYPES ADE

- heritability = 1
- number of traits = 2
- genetic model = A, D, and E
- number of QTNs: A = 2, D = 2, E = 1
- genetic architecture = pleiotropic
- allelic effects = the same used by AlphaSimR

```r
#order in which QTNs where sorted with seed = 10
order_a <- c(1, 2)
order_d <- c(2, 1)
order_e <- c(1)
QTNs_ade <-
  data.frame(
    snp = colnames(QTNs_ade),
    allele = c("A/G", "A/G"),
    chr = c(1, 2),
    pos = c(100, 200),
    cm = c(NA, NA),
    t(QTNs_ade))
custom_a <- list(SP_ade$traits[[1]]@addEff[order_a],
                 SP_ade$traits[[2]]@addEff[order_a])
custom_d <- list(SP_ade$traits[[1]]@domEff[order_d],
                 SP_ade$traits[[2]]@domEff[order_d])
custom_e <- list(SP_ade$traits[[1]]@epiEff[, 3][order_e],
                 SP_ade$traits[[2]]@epiEff[, 3][order_e])
pheno_ade_simple <-  create_phenotypes(
  geno_obj = QTNs_ade,
```

```r
  add_QTN_num = 2,
  dom_QTN_num = 2,
  epi_QTN_num = 1,
  h2 = c(1,1),
  add_effect = custom_a,
  dom_effect = custom_d,
  epi_effect = custom_e,
  ntraits = 2,
  rep = 1,
  vary_QTN = FALSE,
  output_format = "multi-file",
  architecture = "pleiotropic",
  output_dir = "pheno_ade_simple",
  to_r = T,
  sim_method = "custom",
  seed = 2,
  model = "ADE",
  home_dir = getwd(),
  quiet = T)
pheno_ade_simple$Trait_1_H2_1 <- pheno_ade_simple$Trait_1_H2_1 + gv$gv_mu[1]
pheno_ade_simple$Trait_2_H2_1 <- pheno_ade_simple$Trait_2_H2_1 + gv$gv_mu[2]
#Comparison between Trait 1 from AlphaSimR and simplePHENOTYPES
all.equal(pheno_ade_alpha[,1], pheno_ade_simple$Trait_1_H2_1)
# [1] TRUE
#Comparison between Trait 2 from AlphaSimR and simplePHENOTYPES
all.equal(pheno_ade_alpha[,2], pheno_ade_simple$Trait_2_H2_1)
# [1] TRUE
unlink("pheno_ade_simple", recursive = TRUE)
```

#### 1.2 simplePHENOTYPES vs SimPhe

##### 1.2.1 simplePHENOTYPES ADE

- heritability = 1
- number of traits = 2
- genetic model = A, D, and E
- number of QTNs: A = 2, D = 2, E = 1
- genetic architecture = pleiotropic
- allelic effects:
  - A (Trait 1 = 0.3; Trait 2 = 0.1)
  - D (Trait 1 = 0.2; Trait 2 = 0.5)
  - E (Trait 1 = Trait 2 = 0.1)

```r
custom_geometric_a <- list(trait_1 = c(0.3, 0.3),
                           trait_2 = c(0.1, 0.1))
custom_geometric_d <- list(trait_1 = c(0.2, 0.2),
                           trait_2 = c(0.5, 0.5))
```

```r
custom_geometric_e <- list(trait_1 =  0.1,
                           trait_2 =  0.1)
pheno_ade_simple_2 <- create_phenotypes(
  geno_obj = QTNs_ade,
  add_QTN_num = 2,
  dom_QTN_num = 2,
  epi_QTN_num = 1,
  h2 = c(1, 1),
  epi_effect = custom_geometric_e,
  dom_effect = custom_geometric_d,
  add_effect = custom_geometric_a,
  ntraits = 2,
  rep = 1,
  vary_QTN = FALSE,
  output_format = "gemma",
  architecture = "pleiotropic",
  output_dir = "pheno_ade_simple_2",
  seed = 10,
  model = "ADE",
  sim_method = "custom",
  home_dir = getwd(),
  to_r = T,
  quiet = T)
unlink("pheno_ade_simple_2", recursive = TRUE)
```

##### 1.2.2 SimPhe ADE

SimPhe also provides additive and non-additive effects simulation. On the additive effects, both simplePHENOTYPES and SimPhe are equivalent, but they code the major allele in opposite ways (2 = AA in simplePHENOTYPES and 0 = AA in SimPhe). Thus, the allelic effect must have the opposite sign to generate similar results. For instance, in this scenario, the additive allelic effect used on SimPhe was `-0.1` whereas, on simplePHENOTYPES it was `0.1`. The settings for dominance and epistasis were identical.

- Settings used by SimPhe:
- heritability = 1
- number of traits = 2
- genetic model = A, D, and E
- number of QTNs: A = 2, D = 2, E = 1
- mean: trait1 = 0.2 and trait2 = 0.5
- allelic effects:
  - A (Trait 1 = -0.3; Trait 2 = -0.1)
  - D (Trait 1 = 0.2; Trait 2 = 0.5)
  - E (Trait 1 = Trait 2 = 0.1)

```r
snps <- QTNs_ade
rownames(snps) <- snps$snp
```

```
names <- colnames(snps)[-c(1:5)]
snps <- t(snps[,-c(1:5)])
snps <- data.frame(names, snps)
colnames(snps) <- c("IND", "SNP01", "SNP02")
pheno_ade_simphe <- sim.phe(
  sim.pars = "simupars_simphe_ADE.txt",
  fgeno = snps,
  ftype = "snp.head",
  seed = 123,
  fwrite = FALSE)
#Comparison between Trait 1 from simplePHENOTYPES and SimPhe
all.equal(pheno_ade_simple_2$Trait_1_H2_1, pheno_ade_simphe$p1)
# [1] TRUE
#Comparison between Trait 2 from simplePHENOTYPES and SimPhe
all.equal(pheno_ade_simple_2$Trait_2_H2_1, pheno_ade_simphe$p2)
# [1] TRUE
unlink("usedpars.txt")
```

#### 1.3 simplePHENOTYPES vs PhenotypeSimulator

##### 1.3.1 simplePHENOTYPES A

- heritability = 1
- number of traits = 2
- genetic model = A
- number of QTNs: A = 2
- genetic architecture = pleiotropic
- allelic effects: A (QTN 1 = 0.1; QTN 2 = 0.3)

```
custom_geometric_a <- list(trait_1 = c(0.1, 0.3),
                           trait_2 = c(0.1, 0.3))
simplePHENOTYPES_A <- create_phenotypes(
  geno_obj = QTNs_ade,
  add_QTN_num = 2,
  h2 = c(1, 1),
  add_effect = custom_geometric_a,
  ntraits = 2,
  rep = 1,
  vary_QTN = FALSE,
  output_format = "gemma",
  architecture = "pleiotropic",
  output_dir = "simplePHENOTYPES_A",
  seed = 10,
  model = "A",
  sim_method = "custom",
  home_dir = getwd(),
  to_r = T,
```

```
  quiet = T)
unlink("simplePHENOTYPES_A", recursive = TRUE)
```

##### 1.3.2 PhenotypeSimulator A

PhenotypeSimulator provides different options for simulating background and non-genetic effects, but, as far as we know, it does not simulate dominant effects. Thus, we have run a simulation using the geneticFixedEffects() function using the same two QTNs used by simplePHENOTYPES. Results from simplePHENOTYPES and PhenotypeSimulator were also identical. As it is done by simplePHENOTYPES internally, the SNP information was converted from coding 0, 1, and 2 to -1, 0, and 1.

- heritability = 1
- number of traits = 2
- genetic model = A
- number of QTNs: A = 2
- genetic architecture = pleiotropic (pTraitsAffected = 1, pIndependentGenetic = 0, pTraitIndependentGenetic = 0)
- allelic effects: A (QTN 1 = 0.1; QTN 2 = 0.3):

distBeta = "unif"; mBeta = c(0.1, 0.3); sdBeta = c(0.00000000001,0.00000000001)

```
rownames(QTNs_ade) <- QTNs_ade$snp
snps <-
  t(QTNs_ade[c("QTL_2", "QTL_1"),-c(1:5)]) - 1
set.seed(10)
PhenotypeSimulator <- geneticFixedEffects(
  N = 100,
  P = 2,
  X_causal = snps,
  pTraitsAffected = 1,
  pIndependentGenetic = 0,
  pTraitIndependentGenetic = 0,
  distBeta = "unif",
  mBeta = c(0.1, 0.3),
  sdBeta = c(0.00000000001, 0.00000000001))
#Comparison between Trait 1 from simplePHENOTYPES and PhenotypeSimulator
all.equal(simplePHENOTYPES_A$Trait_1_H2_1, as.vector(PhenotypeSimulator$shared[, 1]))
# [1] TRUE
#Comparison between Trait 2 from simplePHENOTYPES and PhenotypeSimulator
all.equal(simplePHENOTYPES_A$Trait_2_H2_1, as.vector(PhenotypeSimulator$shared[, 2]))
# [1] TRUE
```

### 2 Genome-wide association studies (GWAS) on simulated data sets

The following GWAS results were obtained using GEMMA (Zhou and Stephens, 2012, 2014) and figures were generated with the R package CMplot (https://github.com/YinLiLin/R-CMplot).

#### 2.1 Pleiotropy simulation

```
data(SNP55K_maize282_maf04)
custom_effect_a <- list(trait_1 = c(0.2, 0.1, 0.05),
                        trait_2 = c(0.3, 0.2, 0.1))
custom_effect_d <- list(trait_1 = c(0.2),
                        trait_2 = c(0.3))
custom_effect_e <- list(trait_1 = c(0.2, 0.1, 0.05),
                        trait_2 = c(0.3, 0.2, 0.1))

create_phenotypes(
  geno_obj = SNP55K_maize282_maf04,
  add_QTN_num = 3,
  dom_QTN_num = 1,
  h2 = c(0.8, 0.8),
  add_effect = custom_effect_a,
  dom_effect = custom_effect_d,
  ntraits = 2,
  rep = 1,
  output_format = "gemma",
  architecture = "pleiotropic",
  output_dir = "Results_Pleiotropic",
  seed = 100,
  model = "AD",
  sim_method = "custom",
  home_dir = getwd(),
  remove_QTN = F,
  out_geno = "plink",
  quiet = T
)

file.remove("./Results_Pleiotropic/SNP55K_maize282_maf04_noQTN.fam")
file.rename("./Results_Pleiotropic/Simulated_Data__Rep1_Herit_0.8_0.8.fam",
            "./Results_Pleiotropic/SNP55K_maize282_maf04_noQTN.fam")
```

```
include_graphics("Pleiotropy_noQTN.jpg")
```

Table S1: Map and minor allele frequency information of SNPs randomly selected to be used as additive quantitative trait nucleotide in each replication.

| REP | SNP | ALLELE | CHR | POS | CM | MAF |
|---|---|---|---|---|---|---|
| 1 | ss196480696 | G/A | 7 | 151,604,852 | NA | 0.46 |
| 1 | ss196442096 | C/A | 2 | 236,351,458 | NA | 0.44 |
| 1 | ss196509187 | A/G | 10 | 88,613,297 | NA | 0.46 |

Table S2: Map and minor allele frequency information of SNPs randomly selected to be used as dominance quantitative trait nucleotide in each replication.

| REP | SNP | ALLELE | CHR | POS | CM | MAF |
|---|---|---|---|---|---|---|
| 1 | ss196525914 | A/G | 4 | 134,623,817 | NA | 0.41 |

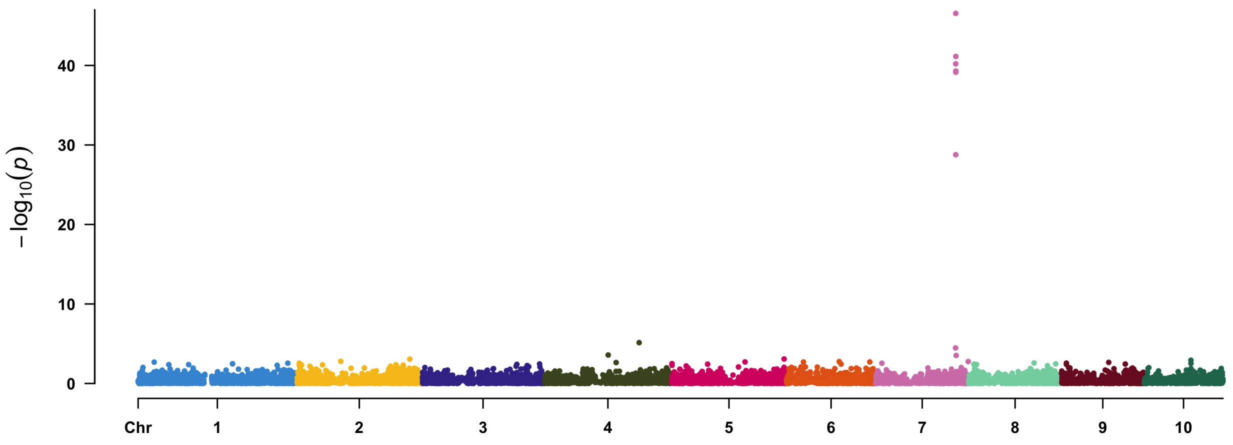


Figure S1: GWAS results obtained from GEMMA using a multi-trait model on a simulated data set with pleiotropic architecture and additive and dominace model.

#### 2.2 Partial Pleiotropy simulation

```
create_phenotypes(
  geno_obj = SNP55K_maize282_maf04,
  ntraits = 2,
  pleio_a = 1,
  pleio_e = 1,
  same_add_dom_QTN = TRUE,
  degree_of_dom = 0.5,
  trait_spec_a_QTN_num = c(2, 2),
  trait_spec_e_QTN_num = c(2, 2),
  h2 = c(0.8, 0.8),
  add_effect = custom_effect_a,
  epi_effect = custom_effect_e,
```

```r
  rep = 1,
  output_dir = "Results_Partially",
  output_format = "gemma",
  architecture = "partially",
  model = "AED",
  seed = 101,
  sim_method = "custom",
  home_dir = getwd(),
  remove_QTN = T,
  quiet = T
)

file.remove("./Results_Partially/SNP55K_maize282_maf04_noQTN.fam")
file.rename("./Results_Partially/Simulated_Data__Rep1_Herit_0.8_0.8.fam",
            "./Results_Partially/SNP55K_maize282_maf04_noQTN.fam")
```

Table S3: Map and minor allele frequency information of SNPs randomly selected to be used as additive and dominance quantitative trait nucleotide in each replication.

| REP | TYPE | TRAIT | SNP | ALLELE | CHR | POS | CM | MAF |
|---|---|---|---|---|---|---|---|---|
| 1 | Pleiotropic | trait_1 | ss196525914 | A/G | 4 | 134,623,817 | NA | 0.41 |
| 1 | trait_specific | trait_1 | ss196418322 | A/G | 4 | 843,215 | NA | 0.41 |
| 1 | trait_specific | trait_1 | ss196504852 | G/A | 1 | 271,913,362 | NA | 0.41 |
| 1 | Pleiotropic | trait_2 | ss196525914 | A/G | 4 | 134,623,817 | NA | 0.41 |
| 1 | trait_specific | trait_2 | ss196476313 | G/A | 7 | 31,144,580 | NA | 0.42 |
| 1 | trait_specific | trait_2 | ss196421720 | A/G | 1 | 277,849,896 | NA | 0.46 |

Table S4: Map and minor allele frequency information of SNPs randomly selected to be used as epistatic quantitative trait nucleotide in each replication.

| REP | TYPE | TRAIT | SNP | ALLELE | CHR | POS | CM | MAF |
|---|---|---|---|---|---|---|---|---|
| 1 | Pleiotropic | trait_1 | ss196501462 | G/A | 1 | 150,112,685 | NA | 0.44 |
| 1 | Pleiotropic | trait_1 | ss196474440 | A/G | 6 | 144,119,580 | NA | 0.43 |
| 1 | trait_specific | trait_1 | ss196504556 | G/A | 3 | 202,038,436 | NA | 0.40 |
| 1 | trait_specific | trait_1 | ss196527150 | A/G | 1 | 4,603,919 | NA | 0.41 |
| 1 | trait_specific | trait_1 | ss196416235 | A/G | 5 | 184,494,302 | NA | 0.44 |
| 1 | trait_specific | trait_1 | ss196428509 | G/A | 1 | 170,057,460 | NA | 0.42 |
| 1 | Pleiotropic | trait_2 | ss196501462 | G/A | 1 | 150,112,685 | NA | 0.44 |
| 1 | Pleiotropic | trait_2 | ss196474440 | A/G | 6 | 144,119,580 | NA | 0.43 |
| 1 | trait_specific | trait_2 | ss196479466 | A/G | 7 | 120,352,276 | NA | 0.44 |
| 1 | trait_specific | trait_2 | ss196431745 | A/G | 1 | 253,493,820 | NA | 0.49 |
| 1 | trait_specific | trait_2 | ss196497469 | C/A | 10 | 74,438,776 | NA | 0.45 |
| 1 | trait_specific | trait_2 | ss196487554 | G/A | 8 | 150,261,503 | NA | 0.45 |

```r
include_graphics("Partial_noQTN.jpg")
```

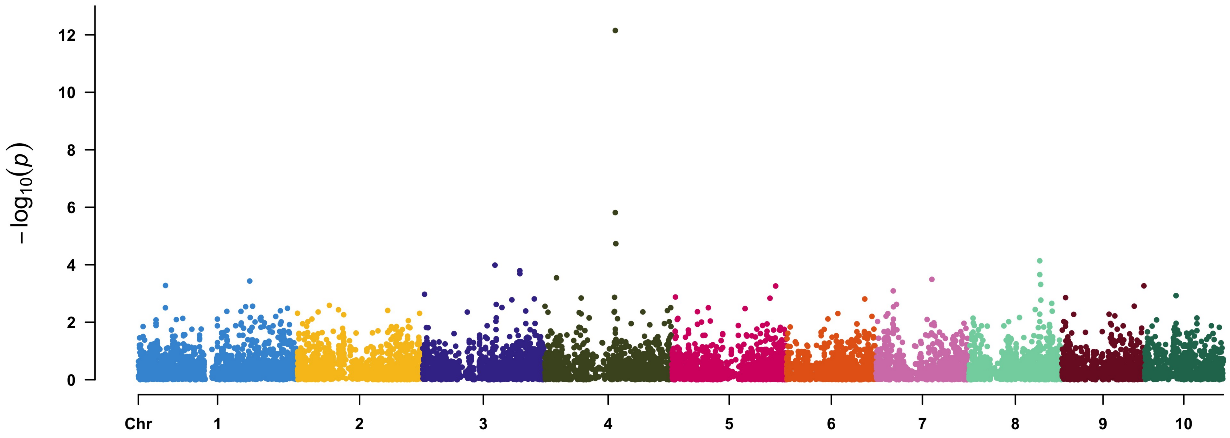


Figure S2: GWAS results obtained from GEMMA using a multi-trait model on a simulated data set with partially pleiotropic architecture and additive and dominace and epistatic model.

### 3 Running a range of settings for an average estimate of time (Figure 3)

#### 3.1 Single Trait simulation

```
data("SNP55K_maize282_maf04")
nn <- 1
st <- list()
file <- list()
setting <- list()
constraints <- list()
for (an in c(0, 0.4)) {
  for (i in c("A", "D", "E", "AD", "AE", "DE", "ADE")) {
    for (j in c(T, F)) {
      if ("D" %in% unlist(strsplit(i,  ""))) {
        constraints$hets <-  "include"
      } else {
        constraints$hets <-  NULL
      }
      setting[[nn]] <- paste("an=", an, "i=", i, "j=", j)
      st[[nn]] <-
        system.time(
          create_phenotypes(
            geno_obj = SNP55K_maize282_maf04,
            add_QTN_num = 2,
            dom_QTN_num = 2,
            epi_QTN_num = 2,
            add_effect = 0.1,
            dom_effect = 0.1,
            epi_effect = 0.1,
            rep = 100,
            h2 = an,
            seed = 100000,
            vary_QTN = j,
            model = i,
            quiet = T,
            output_dir = "ST",
            home_dir = getwd(),
            constraints = constraints
          )
        )
      file[[nn]] <-
        dir("./ST")[grepl("Simulated", dir("./ST"))]
      print(nn)
      nn <- nn + 1
      unlink("ST", recursive = TRUE)
    }
```

```
  }
}
```

#### 3.2 Pleiotropy simulation

```
constraints <- list()
cor_matrix <- matrix(c(1, 0.3, 0.3, 1), 2)
p.time <- list()
big <- c(0.7, 0.3)
mm <- 1
file1 <- list()
setting1 <- list()
for (i in c("A", "D", "E", "AD", "AE", "DE", "ADE")) {
  for (j in c(T, F)) {
    for (x in c("cor_matrix", "NULL")) {
      if ("D" %in% unlist(strsplit(i,  ""))) {
        constraints$hets <-  "include"
      } else {
        constraints$hets <-  NULL
      }
      setting1[[mm]] <-
        paste("i=", i, "j=", j, "x=", x)
      p.time[[mm]] <- system.time(
        create_phenotypes(
          geno_obj  = SNP55K_maize282_maf04,
          add_QTN_num = 2,
          dom_QTN_num = 2,
          epi_QTN_num = 2,
          add_effect = c(0.2, 0.1),
          dom_effect = c(0.2, 0.1),
          epi_effect = c(0.2, 0.1),
          rep = 1000,
          h2 = c(0.7, 0.3),
          seed = 100000,
          vary_QTN = j,
          model = i,
          quiet = TRUE,
          ntraits = 2,
          cor = eval(parse(text = x)),
          architecture = "pleiotropic",
          output_dir = "Pleio",
          home_dir = getwd(),
          constraints = constraints
        )
      )
      file1[[mm]] <-
```

```
        dir("./Pleio")[grepl("Simulated", dir("./Pleio"))]
      print(mm)
      mm <- mm + 1
      unlink("Pleio", recursive = TRUE)
    }
  }
}
```

##### 3.3 Partial Pleiotropy simulation

```
constraints <- list()
cor_matrix <- matrix(c(1, 0.3, 0.3, 1), 2)
file2 <- list()
setting2 <- list()
nn <- 1
pp.time <- list()
for (i in c("A", "D", "E", "AD", "AE", "DE", "ADE")) {
  for (j in c(T, F)) {
    for (x in c("cor_matrix", "NULL")) {
      if ("D" %in% unlist(strsplit(i,  ""))) {
        constraints$hets <-  "include"
      } else {
        constraints$hets <-  NULL
      }
      setting2[[nn]] <-
        paste("i=", i, "j=", j, "x=", x)
      pp.time[[nn]] <-
        system.time(
          create_phenotypes(
            geno_obj = SNP55K_maize282_maf04,
            add_effect = c(0.2, 0.1),
            dom_effect = c(0.2, 0.1),
            epi_effect = c(0.2, 0.1),
            pleio_a = 1,
            pleio_d = 1,
            pleio_e = 1,
            trait_spec_a_QTN_num = c(1, 1),
            trait_spec_d_QTN_num = c(1, 1),
            trait_spec_e_QTN_num = c(1, 1),
            cor = eval(parse(text = x)),
            rep = 1000,
            h2 = c(0.7, 0.3),
            seed = 100000,
            vary_QTN = j,
            model = i,
            quiet = TRUE,
```

```
          ntraits = 2,
          output_dir = "partial",
          architecture = "partially",
          home_dir = getwd(),
          constraints = constraints
        )
      )
    file2[[nn]] <-
      dir("./partial")[grepl("Simulated", dir("./partial"))]
    print(nn)
    nn <- nn + 1
    unlink("partial", recursive = TRUE)
  }
 }
}
```

##### 3.4 Spurious Pleiotropy simulation

```
constraints <- list()
cor_matrix <- matrix(c(1, 0.3, 0.3, 1), 2)
file3 <- list()
setting3 <- list()
nn <- 1
ld.time <- list()
for (i in c("A", "D", "AD")) {
  for (j in c(T, F)) {
    for (x in c("cor_matrix", "NULL")) {
      if ("D" %in% unlist(strsplit(i,  ""))) {
        constraints$hets <-  "include"
      } else {
        constraints$hets <-  NULL
      }
      seed <- 333
      while(length(dir("./LD")[grepl("Simulated", dir("./LD"))]) ==0 ){
      setting3[[nn]] <-
        paste("i=", i, "j=", j, "x=", x)
      ld.time[[nn]] <-
        system.time(
          create_phenotypes(
            geno_obj = SNP55K_maize282_maf04,
            add_QTN_num = 2,
            dom_QTN_num = 2,
            add_effect = c(0.2, 0.1),
            dom_effect = c(0.2, 0.1),
            rep = 1000,
            cor = eval(parse(text = x)),
```

```r
            h2 = c(0.7, 0.3),
            seed = seed,
            vary_QTN = j,
            model = i,
            quiet = TRUE,
            ntraits = 2,
            ld = 0.99,
            output_dir = "LD",
            architecture = "LD",
            home_dir = getwd(),
            type_of_ld = "indirect",
            constraints = constraints
          )
        )
      file3[[nn]] <-
        dir("./LD")[grepl("Simulated", dir("./LD"))]
      print(nn)
      seed <- seed + 11
    }
      unlink("LD", recursive = TRUE)
      nn <- nn + 1
    }
  }
}
```

The script below was used to generate Figure 3. It depicts the time in seconds required by simplePHENOTYPES to simulate each genetic setting.

```r
library(ggplot2)
st <- unlist(lapply(st, function (x) {x["elapsed"]}))
p <- unlist(lapply(p.time, function (x) {x["elapsed"]}))
pp <- unlist(lapply(pp.time, function (x) {x["elapsed"]}))
ld <- unlist(lapply(ld.time, function (x) {x["elapsed"]}))
data <- data.frame(`st`= st, `p`=p, `pp`=pp, `ld`=NA)
data$ld[1:length(ld)] <- ld
data <- reshape2::melt(data)
colnames(data) <- c("Architecture", "Running Time")
#median(data$`Running Time`, na.rm=T)
#2.7905
# median(ld)
# 10.789
# median(p)
# 6.059
# median(pp)
# 15.5725
# median(st)
# 1.164
```

```r
ggplot(data, aes(x=Architecture, y=`Running Time`)) +
  geom_boxplot() + coord_flip() +
  stat_summary(fun.y=mean, geom="point", shape=3, color="black", fill="red") +
  theme(panel.border = element_rect(fill=NA, color="black",
                                    size=0.5, linetype="solid")) +
  scale_x_discrete(name="",
                   limits= c("st", "p", "pp", "ld"),
                   labels=c("Single Trait",
                            "Pleiotropy",
                            "Partial Pleiotropy",
                            "Spurious Pleiotropy")) +
  scale_y_continuous(name="Running Time (s)", breaks= c(0, 25, 50, 75, 100)) +
  theme_bw(base_size = 20) +
  theme( panel.border = element_rect(fill = NA, colour="black")) +
  theme(strip.background =  element_rect(fill = "white", colour = "white")) +
  theme(axis.text.x = element_text(colour = "black", family="Times"),
        axis.ticks = element_blank(), axis.ticks.length = unit(0.1, "cm"))+
  theme(axis.title.y =
          element_text(vjust=1.3, colour = "black", family="Times", size = 25),
        axis.text.y = element_text(colour = "black", family="Times"),
        axis.ticks = element_blank()) +
  theme(panel.grid.major.x = element_blank(), panel.grid.minor = element_blank())
ggsave(file="Figure3.tiff",width=8,height=5,units=c("in"),dpi=300,family="Times")
```
